# Modulating CRISPR gene drive activity through nucleocytoplasmic localization of Cas9 in *S. cerevisiae*

**DOI:** 10.1101/408369

**Authors:** Megan E. Goeckel, Erianna M. Basgall, Isabel C. Lewis, Samantha C. Goetting, Yao Yan, Megan Halloran, Gregory C. Finnigan

**Author notes:** Authors contributed equally (Megan E. Goeckel, Erianna M. Basgall, Isabel C. Lewis, Samantha C. Goetting). Correspondence to: Gregory C. Finnigan.

## Abstract

The bacterial CRISPR/Cas genome editing system has provided a major breakthrough in molecular biology. One use of this technology is within a nuclease-based gene drive. This type of system can install a genetic element within a population at unnatural rates. Combatting of vector-borne diseases carried by metazoans could benefit from a delivery system that bypasses traditional Mendelian laws of segregation. Recently, laboratory studies in fungi, insects, and even mice, have demonstrated successful propagation of CRISPR gene drives and the potential utility of this type of mechanism. However, current gene drives still face challenges including evolved resistance, containment, and the consequences of application in wild populations. In this study, we use an artificial gene drive system in budding yeast to explore mechanisms to modulate nuclease activity of Cas9 through its nucleocytoplasmic localization. We examine non-native nuclear localization sequences on Cas9 fusion proteins *in vivo* and demonstrate that appended signals can titrate gene drive activity and serve as a potential molecular safeguard.

## INTRODUCTION

Control of biological populations is critical to agriculture, ecological conservation, and human health. Numerous methods have been employed to remove invasive species^1,2^, crop-damaging pests^3-6^, or metazoans that harbor diseases^7,8^ including physical barriers, chemical agents, and/or natural predators or competitors. However, the ability to genetically modify an entire species has been hindered by the natural laws of segregation—introduction of a genetic element through natural breeding would require an unattainable number of modified individuals to be released into the wild. However, given the introduction of CRISPR/Cas9 as an efficient, convenient, and universal genome editor^9-15^, a mechanism has been developed that is *Super*-Mendelian in nature: a nuclease “gene drive.”

This arrangement of the CRISPR components is remarkedly simple in design, yet powerful in application. The basic architecture includes a nuclease of choice (usually *S. pyogenes* Cas9, although many alternatives and engineered variants now exist) and the corresponding single guide RNA (sgRNA) integrated within the genome. Placement of Cas9/sgRNA could be at a safe harbor locus, or could delete or disrupt an existing endogenous gene. In the case of the former, the gene drive (GD) would likely also contain a “cargo” gene(s)—the intended genetic element to be delivered to the entire population. This could include any number of variations including endogenous or exogenous DNA to modify the organism itself (e.g. imposed fitness cost) or to aid in the separation between the host and disease-causing agent. Once expressed, the nuclease is primed by the guide RNA to target the *wild-type* copy of the gene (or position) on the homologous chromosome within a diploid genome (within the progeny between the gene drive individual and a wild-type individual) to create a double-stranded break (DSB). The unique arrangement of the GD relative to the DSB allows the expression cassette for Cas9/sgRNA itself to serve as the donor DNA for homology directed repair (HDR). The GD copies itself to the wild-type chromosome to repair the break and replaces the entire endogenous locus; a heterozygous cell (GD/WT) becomes a *homozygous* (GD/GD) cell. Action of a gene drive within a population would allow the rapid “forced” propagation of any genetic element in a small number of generations and would require only a small number of released GD individuals.

There are numerous applications of gene drive biotechnology to control and alter biological populations including global challenges such as eliminating insect-borne diseases^16-18^. Recent experimental^19-24^ and computational studies^25-27^ highlight the potential of GD systems. However, there remain many unknowns surrounding implementation and management of this new technology (including accidental or malicious release of such a system *sans* any safeguard or inhibitory mechanism). Release of a GD-organism has the potential to modify a proportion of the natural population of the chosen species, even using the current available gene drives (for which GD-resistance is still an ongoing issue)^28^. Therefore, it is critical to identify means to control, titrate, inhibit, or reverse gene drive systems to modulate or slow their progression, and as a failsafe should removal of GD individuals become necessary.

Our previous work focused on examination of conserved components of CRISPR gene drives (e.g. nuclease, guide RNA, DNA repair) in budding yeast to identify modes of control, regulation, and inhibition of drive success *in vivo*^29,30^. A variety of molecular mechanisms have been shown to modulate Cas9-based editing including nuclease expression, guide RNA sequence, Cas9-dCas9 fusions, anti-CRISPR mutants, and nucleocytoplasmic shuttling of tagged Cas9. Here, we focused our analysis on appended nuclear localization sequences (NLS) and nuclear export signals (NES) to *S. pyogenes* Cas9-eGFP fusion constructs within an artificial GD system. We tested three nonclassical monopartite NLS sequences, mutated variants, and two NES signals to demonstrate titration of gene drive activity in a diploid yeast model.

## RESULTS

### Non-native nuclear localization signals can direct Cas9 localization *in vivo*

Nucleocytoplasmic transport of macromolecules within eukaryotic cells is highly conserved^31-35^ and has been recognized as a universal requirement of gene editing in living systems—namely, the intended nuclease must gain access to the interior of the nucleus and genomic content. This intracellular trafficking system involves recognition of nuclear import sequences by karyopherins for transit through the nuclear pore complex^36-38^. Use of the CRISPR/Cas system for alteration of the genome typically includes appending one or more NLSs to the nuclease (e.g. Cas9) and the classical NLS^SV40^ is often used for this purpose^10,39,40^. However, several groups have demonstrated that alteration of the nuclear localization of Cas9 can serve as a means to control editing. For instance, the iCas system was constructed to prevent nuclear entry until addition of an external cue^41^. Moreover, design of a split Cas9 included use of a NES sequence to restrict localization of one of the two halves of the nuclease until addition of an exogenous signal^42^. Finally, optimization of CRISPR-based editing in various organisms and cell types has focused on the placement, number, and identity of the included NLS sequence—whether native or non-native to the species of interest^43-49^.

Our previous work in budding yeast demonstrated that the dynamic localization for Cas9 fusions harboring both NLS and NES signals resulted in a variable level of genomic editing (both in haploid and diploid cells)^29^. However, this work focused exclusively on the commonly used NLS^SV40^ signal. A previous study^50^ demonstrated that mutational substitutions to a set of artificially derived NLS signals (from random peptide libraries) allowed for a spectrum of nuclear import efficiencies using a GFP-based reporter system. We sought to test whether alternative nuclear signals could still direct Cas9 to the nucleus and allow for a titration of double-stranded break (DSB) formation in a CRISPR gene drive diploid strain.

Immediately downstream, a modified selectable marker cassette included the highly-expressed and constitutive *CCW12* (cell wall component) promoter sequence driving expression of the *S. pombe HIS5* gene (functional equivalent to yeast *HIS3*). Finally, two (u1) sequences were inserted flanking the entire target locus. To complete action of the gene drive, a high-copy plasmid (pGF-V1220) contained the expression cassette for the guide RNA to prime Cas9 to target the dual (u1) sites was also included within the diploid strain (marked with *LEU2*). The choice to include the guide RNA on a plasmid (rather than placed proximal to the Cas9 gene) was to ensure maximum biosecurity—previous studies have demonstrated that (rapid) separation of the guide and nuclease are a recommended genetic safeguard^21,29^.

Design of our gene drive system allows for the safe and programmable examination of various CRISPR components. Briefly, two sets of “unique” Cas9 target sites (termed u1 and u2)^51^ are positioned flanking both the inducible Cas9-expression cassette (“drive”) and the corresponding locus harboring a selectable marker (“target”). Activation of Cas9 and inclusion of the correct sgRNA fragment (expressed from a plasmid) would cause multiplexing and cleavage of the dual (u1) sites in the target chromosome and repair via homologous recombination using the drive-containing chromosome as the source of donor DNA (Fig. 1). Haploid yeast strains harboring an inducible Cas9-eGFP construct were generated using various C-terminal signals: NLS signals included the classical SV40 and three other nonclassical monopartite signals with sequences containing either four (A), three (B), or two (C) lysine and/or arginine residues within the motif whereas the NES signals (D, E) were derived from the ΦX_3_ΦXb_2_ΦXΦ^52^ consensus (Fig. 2A). Following Cas9 induction in media containing galactose, strains were imaged by fluorescence microscopy (Fig. 2B). For all three non-native NLSs, Cas9 localized to the nucleus (Nup188, a nuclear pore complex component, was marked with mCherry); for both NES-tagged Cas9 fusions, eGFP signal was occluded from the nucleus. However, despite the steady-state localization of the Cas9-eGFP-NES constructs, our previous findings suggested that a small amount of editing (and gene drive activity in diploids) does take place, albeit after significantly longer induction times^29^.

**Figure 1.**
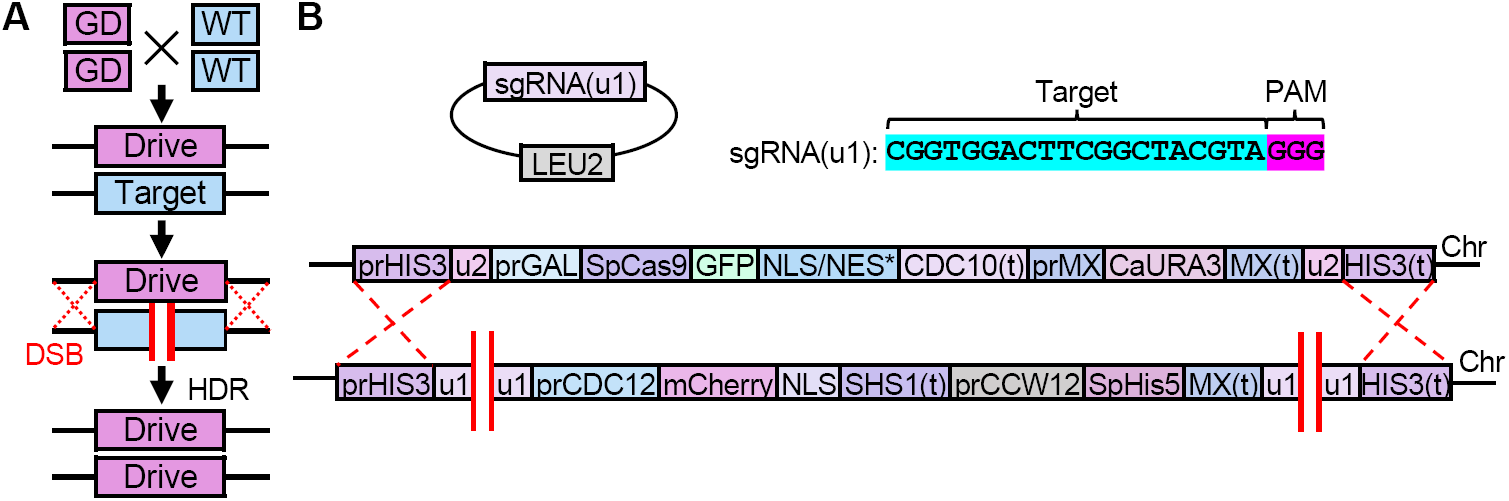
Design of an artificial CRISPR gene drive system in *Saccharomyces cerevisiae*. (A) General schematic of the action of a gene drive in a diploid genome. (B) An artificial gene drive system was constructed at the native yeast *HIS3* locus (*pr*, promoter, *(t)*, terminator sequences). The inducible *GAL1/10* promoter drives expression of the Cas9 nuclease gene (promoter is repressed in the presence of dextrose, and activated in the presence of galactose). A codon-optimized *S. pyogenes* Cas9 contained a C-terminal eGFP fusion followed by a chosen nuclear signal sequence (NLS/NES* see Fig. 2). An inserted terminator (from the yeast *CDC10* gene) was placed downstream of the Cas9 coding sequence followed by a selectable marker cassette. This included the non-native MX-based promoter and terminator sequences driving constitutive expression of the *C. albicans URA3* gene (allowing growth in the absence of uracil). The entire gene drive system was flanked by two identical artificial sites, termed (u2), that do not exist in the yeast genome and provide a convenient genetic safeguard as well as multiplexing at a single locus^29,51^. The design of the gene drive also included an engineered “target” cassette (*bottom*) built within a yeast strain of the opposite mating type at the corresponding *HIS3* locus. This assortment included an artificial “cargo” gene and a yeast-based terminator (from the *SHS1* gene).

**Figure 2.**
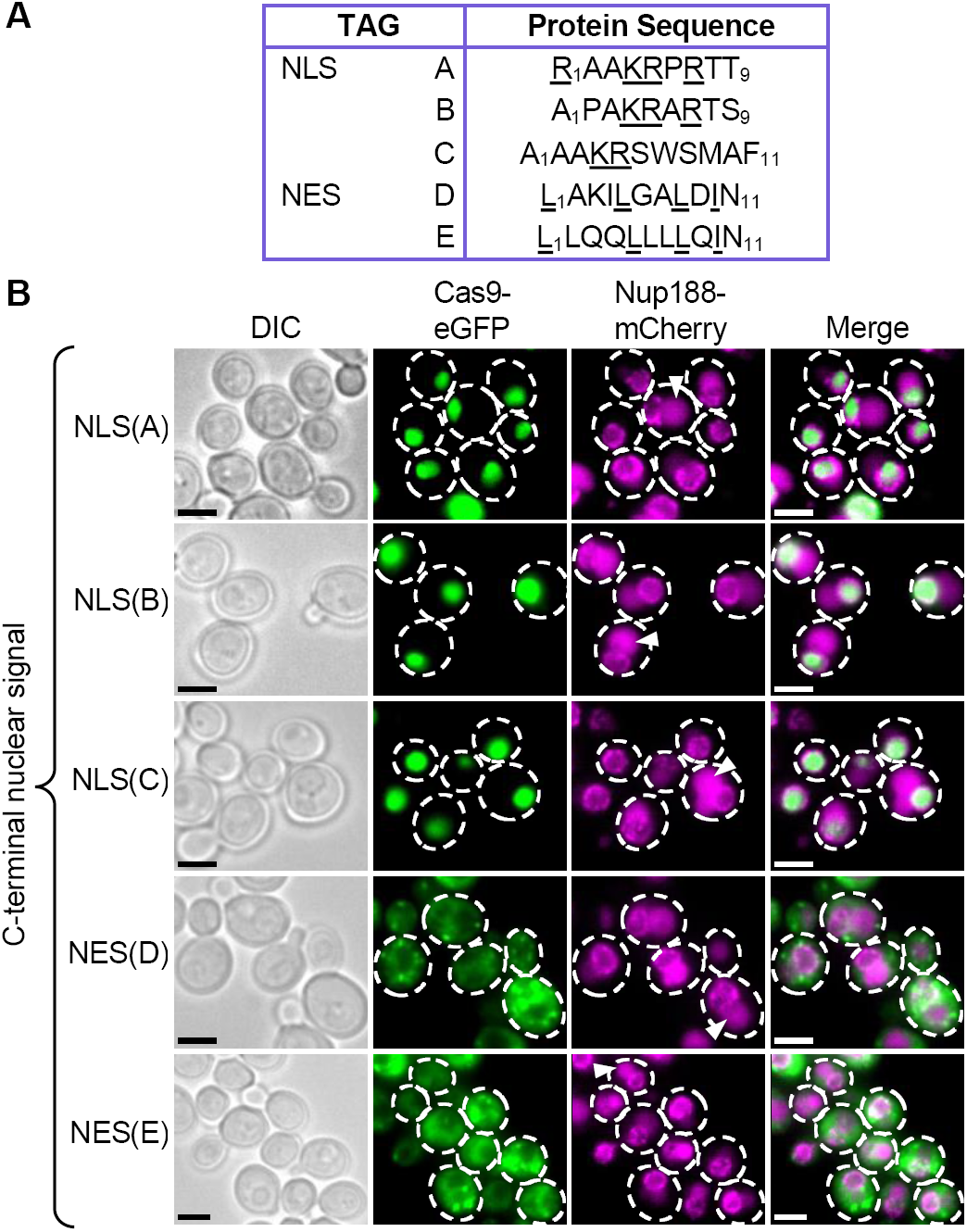
Subcellular localization of *S. pyogenes* Cas9 tagged with various nuclear localization sequences. (A) Table of non-native NLS and NES sequences tested (also see Table 1); basic residues critical to the signal are underlined for NLSs and hydrophobic residues are underlined for NESs. (B) Fluorescence microscopy of live yeast cells containing NLS sequences (GFY-3435 to 3437) or NES sequences (GFY-3438, 3439) fused to eGFP-tagged Cas9. Yeast were cultured in galactose prior to imaging. An integrated copy of Nup188-mCherry marked the nuclear periphery. Representative images are shown; white dotted lines, outline of selected cells. Scale bar, 3 μm. Triangles indicate the yeast vacuole.

### Mutational alterations to NLS or NES sequences can modulate gene drive activity

We tested each NLS and NES sequence appended to Cas9-eGFP for its effect on gene drive activity in diploid yeast. The general methodology employed to assay gene drive function *in vivo* included a variable induction time in galactose followed by recovery on dextrose-containing media (Fig. 3A). Following the formation of single colonies, yeast were transferred to medium lacking histidine (to test for the presence of the *SpHIS5* marker within the target strain). Importantly, our artificial system does not impose any selection for or against action of the gene drive unlike other possible arrangements that challenge cells by selecting for successful DSB/repair events. Compared to the NLS^SV40^-tagged strain, all three artificial NLS signals (A-C) caused a dramatic loss of growth on SD-HIS; constructs harboring the NES (D, E) retained a large number of viable colonies (Fig. 3B). Previous work suggested that mutations to positions along the length of these artificial signals altered their effectiveness at promoting nuclear import by a fluorescence reporter system in live yeast cells^50^. Therefore, we generated twelve substitutions across NLS(A-C) and examined their gene drive activities *in vivo* (Fig. 3C). The total percentage of colonies sensitive to the SD-HIS condition was quantified across multiple trials. As predicted, a NLS^SV40^ tag and a tandem NLS^SV40^-NLS^SV40^ tag allowed for >95% activity at 5 h of Cas9 induction (1, 2). Substitutions to the other classes of NLS (A-C) caused either no change (e.g. 4 and 5) or a moderate decrease in overall drive activity at both time points (e.g. 7, 11, and 14). These results demonstrate that even small alterations to nuclear localization sequences can modify Cas9 editing *in vivo*.

**Table 1.**
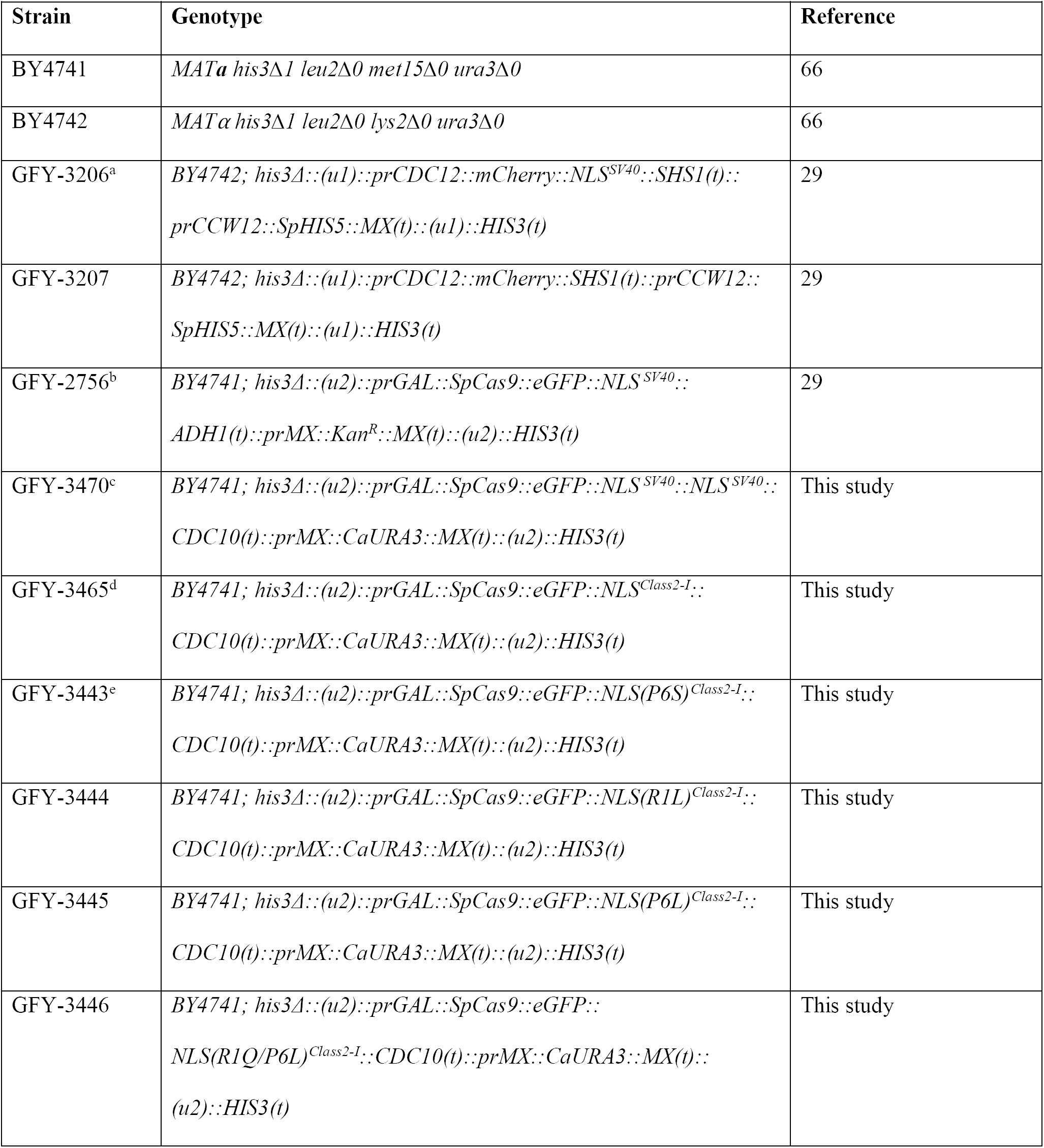
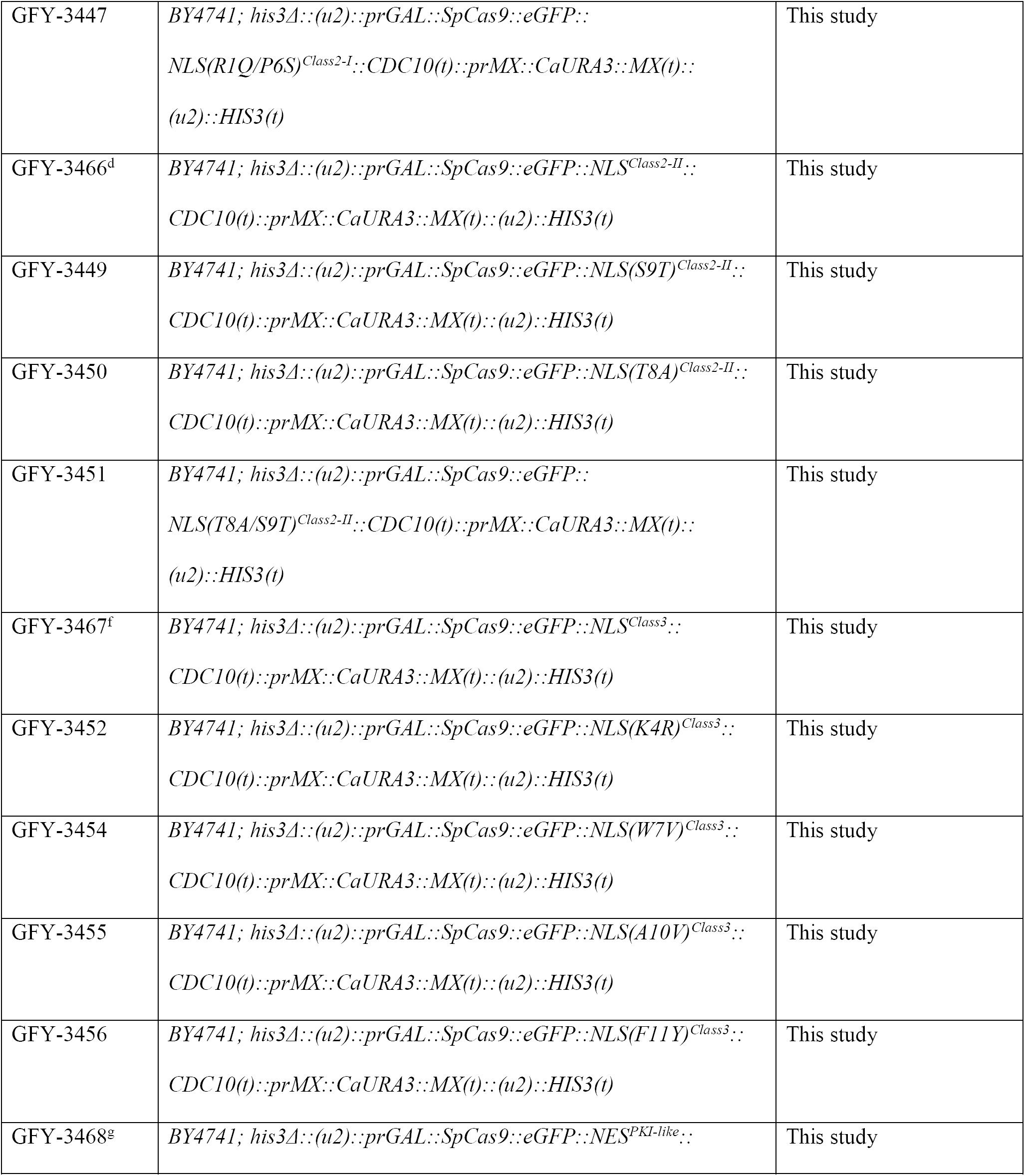
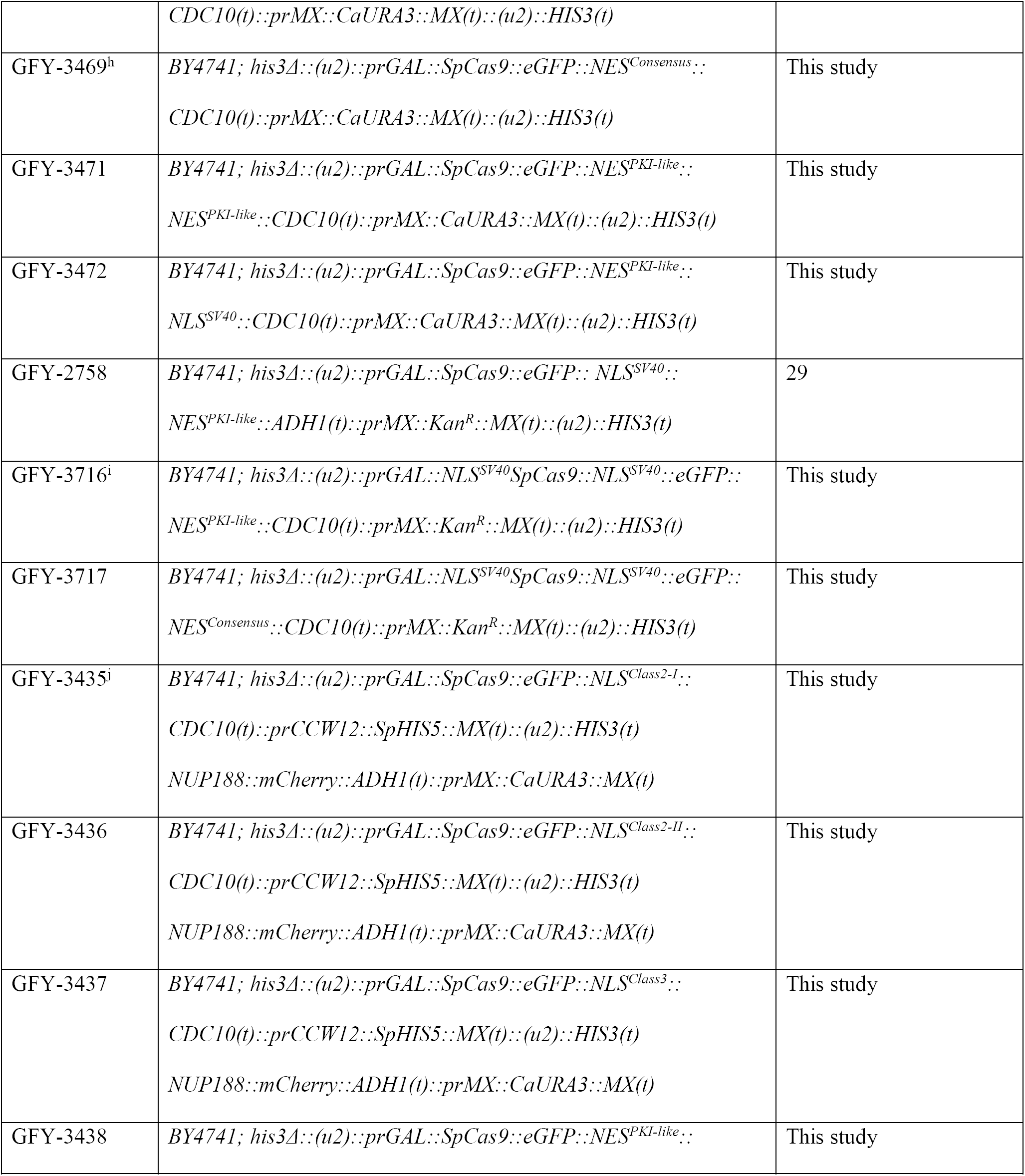
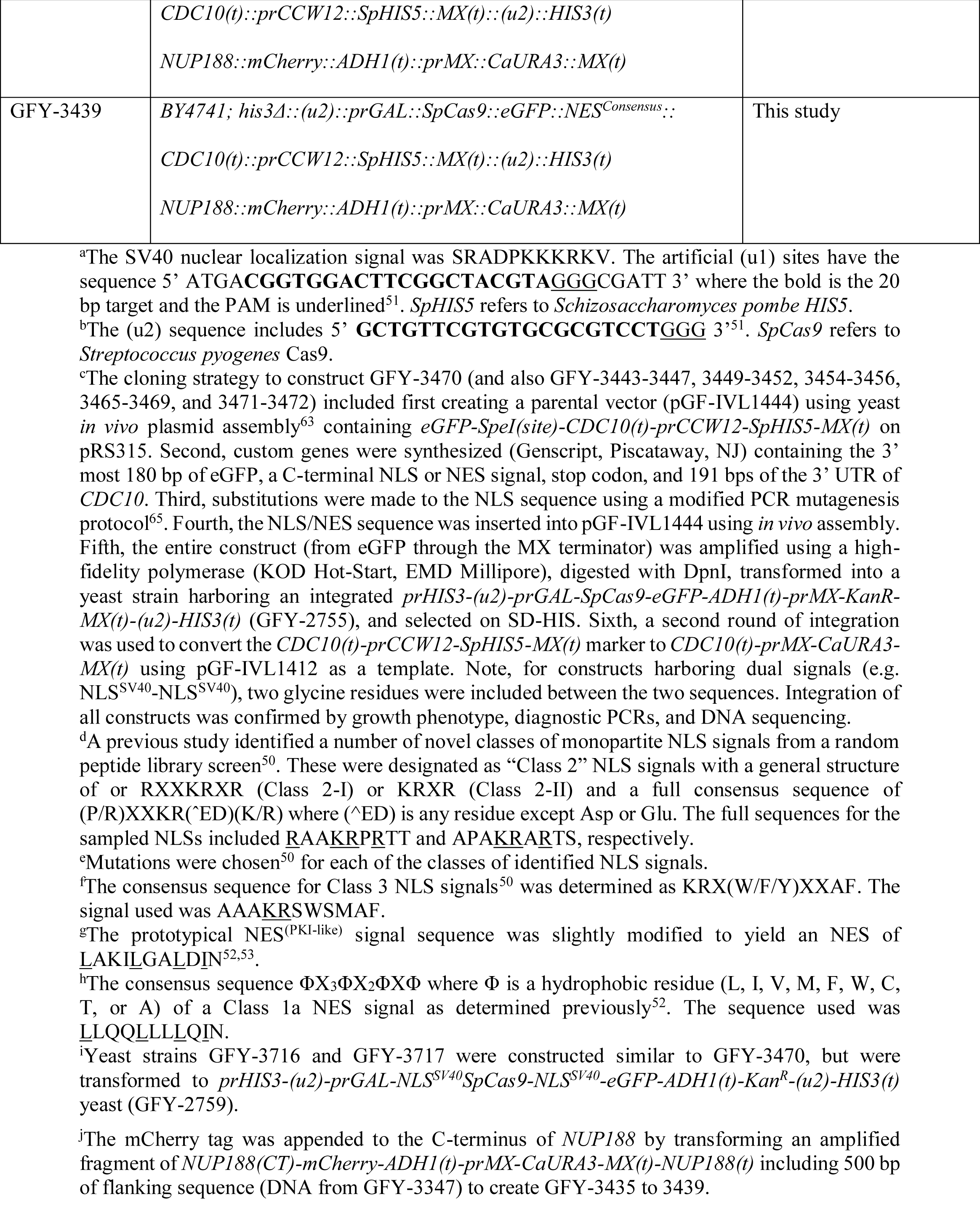
Yeast strains used in this study.

**Figure 3.**
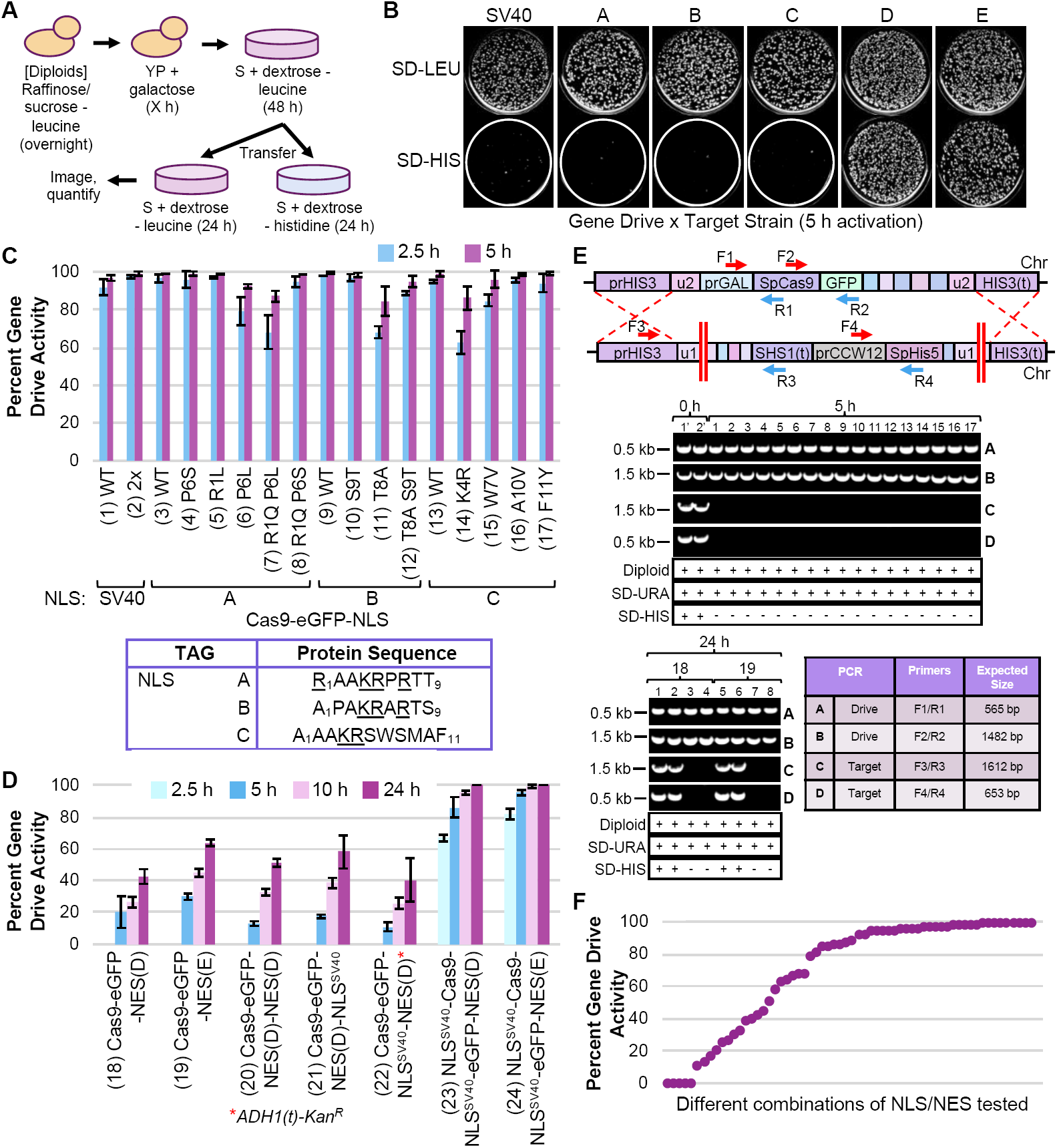
CRISPR gene drives in yeast using various NLS and NES fused to Cas9-eGFP. (A) Schematic of gene drive activation methodology. Following diploid selection (three consecutive rounds), yeast were grown to saturation overnight in media containing raffinose and sucrose lacking leucine. Next, cultures were back-diluted into rich medium containing galactose for a set number of hours, harvested, diluted to approximately 100-500 cells per agar plate (SD-LEU), and incubated for 48 h. Finally, yeast colonies were velvet-transferred to an additional SD-LEU plate and a SD-HIS plate for 24 h before imaging and analysis. (B) Haploid yeast strains (GFY-2756, and GFY-3465-3469) were mated to target yeast strains (GFY-3206 and 3207), diploids were selected, Cas9 was induced for 5 h, and cells were plated as described in (A). (C) Quantifying the efficiency of the gene drive system; the number of colonies sensitive to the SD-HIS condition provided a measure for successful action of the gene drive (referred to as “percent gene drive activity”). Diploid gene drives were tested using strains from (B) and mutational substitutions made to each NLS (GFY-3470, 3443-3447, 3449-3452, and 3454-3456, numbered 1-17) where Cas9 was induced for either 2.5 h or 5 h and quantified for percent gene drive activity. Error, SD. (D) Gene drive strains (GFY-3468, 3469, 3471, 3472, 2758, 3716 and 3717) harboring a NES signal in the absence or presence of additional NLSs were tested for 2.5 h, 5 h, 10 h, and 24 h of Cas9 induction and quantified as in (C). Error, SD. Red asterisk, this construct harbors the *ADH1(t)-prMX-Kan*^*R*^*-MX(t)* cassette following Cas9-eGFP. (E) *Top*, illustration of the general gene drive/target arrangement and the position of oligonucleotides (Supplementary Table S1) used to assess each engineered locus. *Middle*, Diagnostic PCRs were performed on chromosomal DNA from clonal isolates from each gene drive experiment (5 h). The expected band sizes for each PCR (A-D) are shown along with molecular markers. Two isolates were obtained with no galactose activation (1’ and 2’; propagation on only dextrose) to illustrate the presence of both the drive and target loci from GFY-2567 (Strain 1). All colonies were tested for ploidy status (diploid) and growth on SD-URA (drive) and SD-HIS (target). For strain GFY-2756, diploids were tested on G418 media to confirm presence of the drive allele (*Kan*^*R*^) rather than SD-URA. *Below*, A similar analysis of clonal isolates from the NES-containing strains was performed following drive activation (24 h). Two isolates each from (18) and (19) were chosen that were resistant or sensitive to the SD-HIS condition. (F) The gene drive activities from all measured conditions in (C, D) were plotted.

The two NES signals tested on Cas9-eGFP included a PKI-like sequence^53^ as well as a similar NES with added leucine residues present within the motif. A previous study reported that saturation of a NES consensus sequence with additional hydrophobic amino acids or prolines was inhibitory to signal function *in vivo*^52^. Therefore, we tested a NES harboring six total leucine residues as opposed to only three present in the PKI-like motif—the expectation was that a reduction in NES effectiveness would manifest in higher nuclear residence time and gene drive efficiency (Fig. 2A). For any induction times less than 10 h, the gene drive activity (18, 19) remained below 50% (Fig. 3D). We noticed a subtle difference between the effectiveness of the two NES-containing constructs and it appears that the original PKI-like motif (18) provided less drive activity across multiple time points compared to the sequences with added leucine residues (19). A tandem NES^(PKI-like)^-NES^(PKI-like)^ signal (20) did not provide any significant change compared to a single NES alone. Previously, we found that a C-terminal fusion of the dual NLS^(SV40)^-NES^(PKI-like)^ sequence to Cas9-eGFP (22) seemed to phenocopy a fusion of NES^(PKI-like)^ alone^29^. Here, we constructed and tested the reciprocal fusion, NES^(PKI-like)^-NLS^(SV40)^ (21), to determine its effect *in vivo*. We observed that both constructs displayed similar gene drive activities to a single NES signal, yet positioning of the NLS^SV40^ on the extreme C-terminus appeared to provide slightly higher activity (and less contribution from the NES). Finally, we included both NES signals in a construct also harboring two additional NLS^SV40^ sequences—at the N-terminus and fused between Cas9 and eGFP (23, 24). The competition between two NLS sequences versus one NES sequence provided a high level of gene drive activity, with a slight increase in activity from the construct harboring the leucine-rich NES (24). Together, these data illustrate that changes can also be made to either the primary sequence or placement of nuclear export signals to shift the level of editing *in vivo*.

Previous work has demonstrated that gene drives may induce a DSB, but fail to copy the GD cassette to the target chromosome. In such cases, NHEJ causes repair of the broken DNA ends and prevents HR-based propagation of the drive (e.g. GD “resistant” alleles). Therefore, to confirm that yeast sensitive to the SD-HIS condition had lost the target allele at the *HIS3* locus, we obtained clonal isolates from each gene drive experiment, tested for each of the included markers, assayed ploidy status, and examined purified genomic DNA by multiple diagnostic PCRs (Fig. 3E). For diploids lacking galactose treatment (0 h), four independent PCR reactions illustrated the presence of both the gene drive and target constructs. However, for isolates subjected to galactose induction (5 h), only the two PCRs corresponding to the gene drive allele (A, B) were present; reactions for the target allele were unable to amplify the expected fragment (C, D). A similar analysis was performed for eight clonal isolates from the NES strains (18, 19). However, we sampled two isolates that were sensitive and two samples resistant to the SD-HIS condition; diagnostic PCRs illustrated that the target allele was still present for surviving colonies (Fig. 3E, *below*). Finally, we demonstrate a continuum of gene drive activity ranging from 0 to nearly 100% in our yeast model across all Cas9-eGFP fusions with both NLS and NES signals (Fig. 3F).

## DISCUSSION

Given that application of the CRISPR/Cas editing biotechnology in eukaryotic systems requires delivery of Cas9/sgRNA to the nucleus, we focused on methodologies that could provide a new suite of molecular tools to control, inhibit, or modulate gene drive systems *in vivo*, although we recognize these techniques might be used for many types of genomic editing and could apply to alternative uses of the CRISPR system (e.g. dCas9). While there are still few studies on gene drives, the power and potential application for this technology is clear, despite the current challenges and obstacles. The ability to modify an entire population with a genetic element of choice presents numerous advantages including optimizing agricultural crops and animals, prevention of human disease, and ecological control on a large scale. However, ongoing testing of optimal designs including safeguards and controllable drives warrant further research and recent progress has been made in current systems^25,26,54-56^. Previous efforts (both computational and experimental) have highlighted a variety of components that might serve as a platform for control or inhibition of gene drives^21,26,29,30^. Here, we focused on altering the nucleocytoplasmic localization of Cas9 for titration of editing.

While a popular choice for nuclear import across model systems has been the monopartite NLS^SV40^ signal, others have found that varying the position, copy number, or identity of the NLS (native or non-native) can alter genomic editing^43,46,48,57,58^. Given the conservation of nuclear import machinery across eukaryotes and the wide variety of possible nuclear import sequences (native or artificial), this could present a complex platform for tuning or optimizing Cas9 nuclear import in any species of interest. As we have demonstrated here, three artificially generated nonclassical NLS signals allowed for efficient nuclear entry and subsequent gene drive activity *in vivo*; also, some mutational substitutions to the NLS primary sequence partially reduced editing. Future iterations might include modifications to signal positioning (N- or C-terminus, or between fusion proteins), the distance from Cas9, and local residue context surrounding the signal. Of note, our data demonstrates that inclusion of a NES (even when not paired with any NLS signal) still allowed for some level of editing as Cas9 may gain entry to the nucleus through diffusion followed by export; editing *sans* any appended NLS has also been observed in previous studies^29,43^. Therefore, we recommend use of various nuclear signal combinations to modulate and reduce, rather than eliminate, editing. However, nuclear restriction of Cas9 may still be useful when paired with other inhibitory mechanisms such as the AcrIIA2 and AcrIIA4 anti-CRISPR proteins^30^, reduced or programmed expression of nuclease transcript^29^, or other mechanisms to inhibit editing such as degradation of Cas9^59^.

Restriction of Cas9 nuclear localization has also been demonstrated in other cell types through (i) occlusion of a NLS signal or (ii) tethering to the plasma membrane—subsequent release was achieved by activation of a GPCR through addition of an external cue, activation of a protease, and finally, release (via peptide cleavage) of dCas9-NLS and transport into the nucleus^41,60,61^. However, our study provides experimental evidence for use of this general methodology for titration of gene drive activity. We envision that nuclear occlusion of Cas9 could also be achieved by alternative approaches. Expression of an inducible anti-GFP “nanobody”-containing peptide fused with an export signal could provide temporal control to regulate Cas9-GFP nucleocytoplasmic localization. In this way, activity might be reduced at a later point in time (or to a specific portion of a population) by causing an increase in Cas9 nuclear export. A secondary system that could modulate Cas9 activity—inducible by external stimuli—would provide a suite of new options for controlling gene drive propagation within a population. This could be utilized as a molecular safeguard to slow or inhibit drives, allow for the timely use of anti-drives or other countermeasures, or as a means to cycle gene drive-containing organisms with seasonal or environmental changes. Alternatively, modification of the nuclear pore complex might allow for selective entry of a pool of Cas9 fusion constructs while restricting other variants or orthologs. From our analysis, we predict that a combinatorial approach of altering nuclease (transcript and/or protein) levels, NLS/NES signals, cellular traps, and other inducible/tunable systems might be employed in the design of future gene drive systems that are safe, controllable, and/or reversible.

## MATERIALS AND METHODS

### Yeast strains and plasmids

*Saccharomyces cerevisiae* strains used in this study can be found in Table 1. Modern molecular biology techniques were used to generate all engineered constructs^62^. The general strategy included first constructing a *CEN*-based plasmid using *in vivo* assembly in yeast^63^ using a modified lithium acetate transformation protocol^64^. Next, PCR amplified DNA of the entire assembled cassette, followed by treatment with *DpnI* enzyme (to remove template DNA), was integrated into the appropriate haploid yeast genome (for Cas9, the *HIS3* locus). For generation of NLS substitutions, a modified PCR mutagenesis protocol was used^65^. To integrate various eGFP-NLS or eGFP-NES combinations, a second integration construct was generated; universal eGFP and *MX(t)* sequences were used for homologous recombination (Table 1). Finally, selection markers were converted using a third round of integration (e.g. from *SpHIS5* to *CaURA3*) using common *CDC10(t)* and the *MX(t)* sequences. The only plasmid used in this study was the high-copy pRS425-based sgRNA(u1) vector (pGF-V1220)^29^. Diagnostic PCRs and Sanger DNA sequencing of all chromosomal modifications was performed to confirm successful integration events (see Supplementary Fig. S1).

### Culture conditions

Budding yeast were grown on solid agar medium or in liquid cultures; rich media, YPD (2% peptone, 1% yeast extract, 2% dextrose), or synthetic drop-out media (nitrogen base, amino acids, and ammonium sulfate) were used. Synthetic complete media, abbreviated “S.” Prior to galactose metabolism (2%), cultures were grown to saturation in medium containing 2% raffinose and 0.2% sucrose. All sugars were filter sterilized.

### Fluorescence microscopy

Haploid yeast cultures were grown to saturation in synthetic complete medium containing raffinose and sucrose overnight. Next, strains were back-diluted into rich medium containing galactose for 4.5 h at 30°C and prepared on a glass slide^29^. Live cells were examined using a Leica DMI6500 fluorescence microscope (Leica Microsystems, Buffalo Grove, IL). A Leica DFC340 FX camera, 100x lens, and fluorescence filters (Semrock, GFP-4050B-LDKM-ZERO, mCherry-C-LDMK-ZERO) were used. Software for image capture and analysis included Leica Microsystems Application Suite and ImageJ (National Institute of Health). Images were taken using identical exposure times; representative cells were chosen for each image set and scaled together. Scale bars, 3 μm. “Merged” images do not contain any additional processing from the two combined channels.

### CRISPR gene drives in diploid yeast

Haploid yeast strains harboring the *SpCas9* gene drive construct were first transformed with the high-copy *LEU2*-based sgRNA(u1) plasmid (pGF-V1220) and propagated on SD-LEU plates^29^. Next, Cas9/sgRNA-containing haploids were mated to target yeast strains of the opposite mating type on rich media for 24 h and transferred to diploid selection plates (SD-LEU-HIS) for three consecutive stages of selection. Diploids were cultured overnight to saturation (raffinose/sucrose), grown in rich medium containing galactose for (0 to 24 h) to express Cas9, diluted, spread onto SD-LEU medium at a density of 100-500 cells per plate, and incubated at 30°C for 48 h. Finally, colonies were replica-plated by velvet transfer to a second SD-LEU and SD-HIS plate for 18-24 h, imaged, and the total surviving colony number was quantified in a single-blind fashion (between 100-200 colonies counted for each condition). Gene drive and target genome status were interrogated by diagnostic PCRs (see Supplementary Fig. S2) on isolated chromosomal DNA (also confirmed as diploids^29^). Genetic safeguards to contain yeast gene drives included the use of artificial sequences^51^ programmed at the *HIS3* locus (u1 sites for targeting), a self-excision module (u2 sites) flanking all Cas9 constructs^29^, an inducible promoter driving Cas9 expression, and sgRNA plasmids on an unstable high-copy vector^21,29^.

## ETHICS APPROVAL AND CONSENT TO PARTICIPATE

Not applicable.

## CONSENT FOR PUBLICATION

Not applicable.

## AVAILABILITY OF DATA AND MATERIALS

The datasets used and analyzed during the current study are available from the corresponding author on reasonable request.

## AUTHOR CONTRIBUTIONS

M.E.G., E.M.B, I.C.L., S.C.G., Y.Y., M.H., and G.C.F built all reagents (plasmids and yeast) and performed all experiments. M.E.G., E.M.B, I.C.L., S.C.G, and G.C.F. performed all data analyses and figure preparation. G.C.F. wrote the manuscript.

## ACKNOWLEDGEMENTS

We thank Rachael Giersch, Emily Roggenkamp, and Madison Schrock (Kansas State University) for useful comments and laboratory assistance.

## ABBREVIATIONS

CRISPR: clustered regularly interspaced short palindromic repeats
GD: gene drive
DSB: double stranded break
HR: homologous recombination
HDR: homology directed repair
NHEJ: non-homologous end joining
sgRNA: single guide RNA fragment
NLS: nuclear localization sequence
NES: nuclear export signal
GPCR: G-protein coupled receptor
DIC: differential interference contrast.

## REFERENCES

1 Bajwa, A. A. et al. Biology and management of two important Conyza weeds: a global review. Environmental science and pollution research international 23, 24694-24710, doi:10.1007/s11356-016-7794-7 (2016).

2 Neilson, B. J., Wall, C. B., Mancini, F. T. & Gewecke, C. A. Herbivore biocontrol and manual removal successfully reduce invasive macroalgae on coral reefs. PeerJ 6, e5332, doi:10.7717/peerj.5332 (2018).

3 Kergunteuil, A. et al. Biological Control beneath the Feet: A Review of Crop Protection against Insect Root Herbivores. Insects 7, DOI:10.3390/insects7040070 (2016).

4 Khan, Z., Midega, C. A., Hooper, A. & Pickett, J. Push-Pull: Chemical Ecology-Based Integrated Pest Management Technology. Journal of chemical ecology 42, 689-697, doi:10.1007/s10886-016-0730-y (2016).

5 Stenberg, J. A., Heil, M., Ahman, I. & Bjorkman, C. Optimizing Crops for Biocontrol of Pests and Disease. Trends in plant science 20, 698-712, doi:10.1016/j.tplants.2015.08.007 (2015).

6 Peterson, J. A., Ode, P. J., Oliveira-Hofman, C. & Harwood, J. D. Integration of Plant Defense Traits with Biological Control of Arthropod Pests: Challenges and Opportunities. Frontiers in plant science 7, 1794, doi:10.3389/fpls.2016.01794 (2016).

7 Delves, M. J., Angrisano, F. & Blagborough, A. M. Antimalarial Transmission-Blocking Interventions: Past, Present, and Future. Trends in parasitology, DOI:10.1016/j.pt.2018.07.001 (2018).

8 Tizifa, T. A. et al. Prevention Efforts for Malaria. Current tropical medicine reports 5, 41-50, doi:10.1007/s40475-018-0133-y (2018).

9 Jinek, M. et al. A programmable dual-RNA-guided DNA endonuclease in adaptive bacterial immunity. Science (New York, N.Y.) 337, 816-821, doi:10.1126/science.1225829 (2012).

10 Jinek, M. et al. RNA-programmed genome editing in human cells. eLife 2, e00471, doi:10.7554/eLife.00471 (2013).

11 Sternberg, S. H., Redding, S., Jinek, M., Greene, E. C. & Doudna, J. A. DNA interrogation by the CRISPR RNA-guided endonuclease Cas9. Nature 507, 62-67, doi:10.1038/nature13011 (2014).

12 Doudna, J. A. & Charpentier, E. Genome editing. The new frontier of genome engineering with CRISPR-Cas9. Science (New York, N.Y.) 346, 1258096, doi:10.1126/science.1258096 (2014).

13 Estrela, R. & Cate, J. H. Energy biotechnology in the CRISPR-Cas9 era. Current opinion in biotechnology 38, 79-84, doi:10.1016/j.copbio.2016.01.005 (2016).

14 Sternberg, S. H. & Doudna, J. A. Expanding the Biologist’s Toolkit with CRISPR-Cas9. Molecular cell 58, 568-574, doi:10.1016/j.molcel.2015.02.032 (2015).

15 Wright, A. V., Nunez, J. K. & Doudna, J. A. Biology and Applications of CRISPR Systems: Harnessing Nature’s Toolbox for Genome Engineering. Cell 164, 29-44, doi:10.1016/j.cell.2015.12.035 (2016).

16 Esvelt, K. M. & Gemmell, N. J. Conservation demands safe gene drive. PLoS biology 15, e2003850, doi:10.1371/journal.pbio.2003850 (2017).

17 Prowse, T. A. A. et al. Dodging silver bullets: good CRISPR gene-drive design is critical for eradicating exotic vertebrates. Proceedings. Biological sciences 284, DOI:10.1098/rspb.2017.0799 (2017).

18 Courtier-Orgogozo, V., Morizot, B. & Boete, C. Using CRISPR-based gene drive for agriculture pest control. EMBO reports, DOI:10.15252/embr.201744822 (2017).

19 Roggenkamp, E. et al. CRISPR-UnLOCK: multipurpose Cas9-based strategies for Conversion of yeast libraries and strains. Frontiers in microbiology 8, 1773, doi:10.3389/fmicb.2017.01773 (2017).

20 Shapiro, R. S. et al. A CRISPR-Cas9-based gene drive platform for genetic interaction analysis in Candida albicans. Nature microbiology 3, 73-82, doi:10.1038/s41564-017-0043-0 (2018).

21 DiCarlo, J. E., Chavez, A., Dietz, S. L., Esvelt, K. M. & Church, G. M. Safeguarding CRISPR-Cas9 gene drives in yeast. Nature biotechnology 33, 1250-1255, doi:10.1038/nbt.3412 (2015).

22 Gantz, V. M. et al. Highly efficient Cas9-mediated gene drive for population modification of the malaria vector mosquito Anopheles stephensi. Proceedings of the National Academy of Sciences of the United States of America 112, E6736-6743, doi:10.1073/pnas.1521077112 (2015).

23 Hammond, A. et al. A CRISPR-Cas9 gene drive system targeting female reproduction in the malaria mosquito vector Anopheles gambiae. Nature biotechnology 34, 78-83, doi:10.1038/nbt.3439 (2016).

24 Grunwald, H. A. et al. Super-Mendelian inheritance mediated by CRISPR/Cas9 in the female mouse germline. bioRxiv, DOI:10.1101/362558 (2018).

25 Noble, C., Olejarz, J., Esvelt, K. M., Church, G. M. & Nowak, M. A. Evolutionary dynamics of CRISPR gene drives. Science advances 3, e1601964, doi:10.1126/sciadv.1601964 (2017).

26 Vella, M. R., Gunning, C. E., Lloyd, A. L. & Gould, F. Evaluating strategies for reversing CRISPR-Cas9 gene drives. Scientific reports 7, 11038, doi:10.1038/s41598-017-10633-2 (2017).

27 Dhole, S., Vella, M. R., Lloyd, A. L. & Gould, F. Invasion and migration of spatially self-limiting gene drives: A comparative analysis. Evolutionary applications 11, 794-808, doi:10.1111/eva.12583 (2018).

28 Noble, C., Adlam, B., Church, G. M., Esvelt, K. M. & Nowak, M. A. Current CRISPR gene drive systems are likely to be highly invasive in wild populations. eLife 7, DOI:10.7554/eLife.33423 (2018).

29 Roggenkamp, E. et al. Tuning CRISPR-Cas9 Gene Drives in Saccharomyces cerevisiae. G3 (Bethesda, Md.) 8, 999-1018, doi:10.1534/g3.117.300557 (2018).

30 Basgall, E. M. et al. Gene drive inhibition by the anti-CRISPR proteins AcrIIA2 and AcrIIA4 in Saccharomyces cerevisiae. Microbiology (Reading, England) 164, 464-474, doi:10.1099/mic.0.000635 (2018).

31 Kim, Y. H., Han, M. E. & Oh, S. O. The molecular mechanism for nuclear transport and its application. Anatomy & cell biology 50, 77-85, doi:10.5115/acb.2017.50.2.77 (2017).

32 Bauer, N. C., Doetsch, P. W. & Corbett, A. H. Mechanisms Regulating Protein Localization. Traffic (Copenhagen, Denmark) 16, 1039-1061, doi:10.1111/tra.12310 (2015).

33 Sorokin, A. V., Kim, E. R. & Ovchinnikov, L. P. Nucleocytoplasmic transport of proteins. Biochemistry. Biokhimiia 72, 1439–1457 (2007).

34 Poon, I. K. & Jans, D. A. Regulation of nuclear transport: central role in development and transformation? Traffic (Copenhagen, Denmark) 6, 173-186, doi:10.1111/j.1600-0854.2005.00268.x (2005).

35 Stewart, M. Molecular mechanism of the nuclear protein import cycle. Nature reviews. Molecular cell biology 8, 195-208, doi:10.1038/nrm2114 (2007).

36 DeGrasse, J. A. et al. Evidence for a shared nuclear pore complex architecture that is conserved from the last common eukaryotic ancestor. Molecular & cellular proteomics: MCP 8, 2119-2130, doi:10.1074/mcp.M900038-MCP200 (2009).

37 Wente, S. R. & Rout, M. P. The nuclear pore complex and nuclear transport. Cold Spring Harbor perspectives in biology 2, a000562, doi:10.1101/cshperspect.a000562 (2010).

38 O’Reilly, A. J., Dacks, J. B. & Field, M. C. Evolution of the karyopherin-beta family of nucleocytoplasmic transport factors; ancient origins and continued specialization. PloS one 6, e19308, doi:10.1371/journal.pone.0019308 (2011).

39 Glass, Z., Lee, M., Li, Y. & Xu, Q. Engineering the Delivery System for CRISPR-Based Genome Editing. Trends in biotechnology 36, 173-185, doi:10.1016/j.tibtech.2017.11.006 (2018).

40 Kalderon, D., Roberts, B. L., Richardson, W. D. & Smith, A. E. A short amino acid sequence able to specify nuclear location. Cell 39, 499–509 (1984).

41 Liu, K. I. et al. A chemical-inducible CRISPR-Cas9 system for rapid control of genome editing. Nature chemical biology 12, 980-987, doi:10.1038/nchembio.2179 (2016).

42 Zetsche, B., Volz, S. E. & Zhang, F. A split-Cas9 architecture for inducible genome editing and transcription modulation. Nature biotechnology 33, 139-142, doi:10.1038/nbt.3149 (2015).

43 Hu, P., Zhao, X., Zhang, Q., Li, W. & Zu, Y. Comparison of Various Nuclear Localization Signal-Fused Cas9 Proteins and Cas9 mRNA for Genome Editing in Zebrafish. G3 (Bethesda, Md.) 8, 823-831, doi:10.1534/g3.117.300359 (2018).

44 Menoret, S. et al. Homology-directed repair in rodent zygotes using Cas9 and TALEN engineered proteins. Scientific reports 5, 14410, doi:10.1038/srep14410 (2015).

45 Torres-Ruiz, R. et al. Efficient Recreation of t(11;22) EWSR1-FLI1+ in Human Stem Cells Using CRISPR/Cas9. Stem cell reports 8, 1408-1420, doi:10.1016/j.stemcr.2017.04.014 (2017).

46 Staahl, B. T. et al. Efficient genome editing in the mouse brain by local delivery of engineered Cas9 ribonucleoprotein complexes. Nature biotechnology 35, 431-434, doi:10.1038/nbt.3806 (2017).

47 Weninger, A., Glieder, A. & Vogl, T. A toolbox of endogenous and heterologous nuclear localization sequences for the methylotrophic yeast Pichia pastoris. FEMS yeast research 15, DOI:10.1093/femsyr/fov082 (2015).

48 Cong, L. et al. Multiplex genome engineering using CRISPR/Cas systems. Science (New York, N.Y.) 339, 819-823, doi:10.1126/science.1231143 (2013).

49 Shen, B. et al. Generation of gene-modified mice via Cas9/RNA-mediated gene targeting. Cell research 23, 720-723, doi:10.1038/cr.2013.46 (2013).

50 Kosugi, S. et al. Six classes of nuclear localization signals specific to different binding grooves of importin alpha. The Journal of biological chemistry 284, 478-485, doi:10.1074/jbc.M807017200 (2009).

51 Finnigan, G. C. & Thorner, J. mCAL: a new approach for versatile multiplex action of Cas9 using one sgRNA and loci flanked by a programmed target sequence. G3 (Bethesda, Md.) 6, 2147-2156, doi:10.1534/g3.116.029801 (2016).

52 Kosugi, S., Hasebe, M., Tomita, M. & Yanagawa, H. Nuclear export signal consensus sequences defined using a localization-based yeast selection system. Traffic (Copenhagen, Denmark) 9, 2053-2062, doi:10.1111/j.1600-0854.2008.00825.x (2008).

53 Wen, W., Meinkoth, J. L., Tsien, R. Y. & Taylor, S. S. Identification of a signal for rapid export of proteins from the nucleus. Cell 82, 463–473 (1995).

54 Hammond, A. M. et al. The creation and selection of mutations resistant to a gene drive over multiple generations in the malaria mosquito. PLoS genetics 13, e1007039, doi:10.1371/journal.pgen.1007039 (2017).

55 Drury, D. W., Dapper, A. L., Siniard, D. J., Zentner, G. E. & Wade, M. J. CRISPR/Cas9 gene drives in genetically variable and nonrandomly mating wild populations. Science advances 3, e1601910, doi:10.1126/sciadv.1601910 (2017).

56 Champer, J. et al. Reducing resistance allele formation in CRISPR gene drive. Proceedings of the National Academy of Sciences of the United States of America, DOI:10.1073/pnas.1720354115 (2018).

57 Wang, Y. et al. A ‘suicide’ CRISPR-Cas9 system to promote gene deletion and restoration by electroporation in Cryptococcus neoformans. Scientific reports 6, 31145, doi:10.1038/srep31145 (2016).

58 Osakabe, Y. et al. Optimization of CRISPR/Cas9 genome editing to modify abiotic stress responses in plants. Scientific reports 6, 26685, doi:10.1038/srep26685 (2016).

59 Maji, B. et al. Multidimensional chemical control of CRISPR-Cas9. Nature chemical biology 13, 9-11, doi:10.1038/nchembio.2224 (2017).

60 Kipniss, N. H. et al. Engineering cell sensing and responses using a GPCR-coupled CRISPR-Cas system. Nature communications 8, 2212, doi:10.1038/s41467-017-02075-1 (2017).

61 Baeumler, T. A., Ahmed, A. A. & Fulga, T. A. Engineering Synthetic Signaling Pathways with Programmable dCas9-Based Chimeric Receptors. Cell reports 20, 2639-2653, doi:10.1016/j.celrep.2017.08.044 (2017).

62 Sambrook, J. & Russell, D. W. Molecular Cloning: A Laboratory Manual. 3rd edn, (Cold Spring Harbor Laboratory Press, 2001).

63 Finnigan, G. C. & Thorner, J. Complex *in vivo* ligation using homologous recombination and high-efficiency plasmid rescue from *Saccharomyces cerevisiae*. Bio-protocol 5, e1521. http://www.bio-protocol.org/e1521 (2015).

64 Eckert-Boulet, N., Pedersen, M. L., Krogh, B. O. & Lisby, M. Optimization of ordered plasmid assembly by gap repair in Saccharomyces cerevisiae. Yeast (Chichester, England) 29, 323-334, doi:10.1002/yea.2912 (2012).

65 Zheng, L., Baumann, U. & Reymond, J. L. An efficient one-step site-directed and site-saturation mutagenesis protocol. Nucleic acids research 32, e115, doi:10.1093/nar/gnh110 (2004).

66 Brachmann, C. B. et al. Designer deletion strains derived from Saccharomyces cerevisiae S288C: a useful set of strains and plasmids for PCR-mediated gene disruption and other applications. Yeast (Chichester, England) 14, 115-132, doi:10.1002/(sici)1097-0061(19980130)14:2<115::aid-yea204>3.0.co;2-2 (1998).

